# Hyperactivation of MEK1 in cortical glutamatergic neurons results in projection axon deficits and aberrant motor learning

**DOI:** 10.1101/2023.11.06.565901

**Authors:** George R. Bjorklund, Katherina P. Rees, Kavya Balasubramanian, Lauren T. Hewitt, Kenji Nishimura, Jason M. Newbern

## Abstract

Abnormal Extracellular Regulated Kinase 1/2 (ERK1/2) signaling is linked to multiple neurodevelopmental diseases, especially the RASopathies, which typically exhibit ERK1/2 hyperactivation in neurons and non-neuronal cells. To better understand how excitatory neuron-autonomous ERK1/2 activity regulates the development of the mouse motor cortex, we conditionally expressed a hyperactive MEK1^S217/221E^ variant using *Nex/NeuroD6:Cre*. Our results show that MEK1^S217/221E^ expression led to persistent hyperactivation of ERK1/2 in neocortical axons, but not excitatory neuron somas or nuclei. We noted reduced axonal arborization of multiple subcortical target domains in mutants and reduced cortical expression of the activity dependent gene, ARC. These changes did not coincide with significant differences in voluntary locomotor activity or motor performance in the accelerating rotarod task. However, motor learning in a single-pellet retrieval task was significantly diminished in *Nex/NeuroD6:Cre*; *MEK1^S217/221E^* mutants. Restriction of MEK1^S217/221E^ expression to layer V cortical neurons recapitulated axonal outgrowth deficits, however, had no effect on motor learning. Collectively, these results indicate that within the cortex, glutamatergic neuron-autonomous hyperactivation of MEK1 is sufficient to drive deficits in axon outgrowth, activity dependent gene expression, and skilled motor learning.

**Summary statement:** MEK-ERK1/2 hyperactivation in developing cortical excitatory neurons is sufficient to decrease long-range axonal outgrowth, which coincides with reduced Arc expression and deficits in aspects of skilled motor learning by adulthood.

## Introduction

Many extracellular cues important for forebrain development, such as fibroblast growth factors (FGFs) and neurotrophins, activate receptor tyrosine kinases (RTKs) to modulate a wide range of cellular behaviors. Ligand-bound RTKs initiate intracellular signaling via numerous kinase cascades, including the PI3K, PLC, PKC, and Extracellular Regulated Kinase 1/2 (ERK1/2) pathways (Lemmon and Schlessinger, 2010). Activation of ERK1/2 downstream of RTK signaling has been well-studied, which requires RAS-dependent activation of RAF isoforms and subsequent phosphorylation of MEK1 and MEK2 at two conserved residues in the kinase domain, Ser^218/222^ and Ser^222/226^, respectively (Klomp et al., 2021; Lavoie et al., 2020). Activated MEK1/2 then phosphorylates ERK1 and ERK2, which ultimately act upon hundreds of cytosolic and nuclear substrates (Ünal et al., 2017). ERK1/2 signaling has frequently been linked to the control of cell proliferation, differentiation, survival, synaptic plasticity, and learning (Atkins et al., 2001; Gómez-Palacio-Schjetnan and Escobar, 2008; Jeanneteau et al., 2010; Kalil and Dent, 2014; Mebratu and Tesfaigzi, 2009; Nowaczyk et al., 2014; Polleux and Snider, 2010; Thomas and Huganir, 2004). However, this broad repertoire obscures the intricate, context-dependent functions of ERK1/2 signaling that are highly contingent on the precise developmental stage, cell type, and/or extracellular cue (Santini et al., 2019; Shaul and Seger, 2007). Developing neurons in particular exhibit complex, subtype specific responses to modifications in ERK1/2 activity that are not fully understood (Ryu and Lee, 2016).

Key regulators in the RTK-ERK1/2 signaling network are frequently mutated in a family of developmental syndromes known collectively as the “RASopathies” (Jindal et al., 2015; Kim and Baek, 2019; Rauen, 2013; Rauen, 2022). Well-defined mutations have been identified in individual syndromes within the RASopathy superfamily, such as Noonan syndrome (NS), Neurofibromatosis type 1 (NF1), Costello syndrome (CS), and Cardio-facio-cutaneous syndrome (CFC). NS and NF1 are common RASopathies with a combined frequency of ∼1:2000 births (Rauen, 2013; Tidyman and Rauen, 2009). However, >50 distinct RASopathy mutations in over 10 different genes within the RTK-ERK1/2 network have been reported (Brown et al., 2012; Gutmann et al., 2012; Jindal et al., 2015; Kaul et al., 2015; Rauen, 2022). Upstream RASopathy mutations in NF1, SHP2, and RAS have been shown to modulate PI3K/Akt, PKC, and/or Rho/ROCK activity, in addition to the ERK1/2 pathway (Anastasaki and Gutmann, 2014; Brown et al., 2012; Castellano and Downward, 2011; Kaul et al., 2015). ERK1/2 hyperactivation is, however, thought to be a core, common feature of nearly all RASopathies. Thus, pharmacological inhibitors of ERK1/2 regulators have been an important focus for therapeutic development (Guilding et al., 2007; López-Juárez et al., 2017; Nakamura et al., 2009; Wang et al., 2012; Wu et al., 2011). ERK1/2 also represents a convergent target downstream of multiple genetic risk factors for autism and is abnormally activated in models of Fragile X and 16p11.2 deletion syndrome (Chung et al., 2021; Osterweil et al., 2013; Wen et al., 2016). Therefore, understanding the cell-specific effects of ERK1/2 activity in the developing forebrain may shed light on potential neuropathological processes in multiple developmental disorders (Bernardino et al., 2022; Li et al., 2005; Roskoski, 2019).

MRI-based studies suggest that differences in both cortical structural and functional connectivity are a critical aspect of intellectual disability and cognitive changes in RASopathies and autism (Aydin et al., 2016; Chang et al., 2016; Fattah et al., 2021; Filippi et al., 2013; Ibrahim et al., 2017; Owen et al., 2018; Tomson et al., 2015; Zamboni et al., 2007). Changes in motor cortex function have been detected in RASopathies and individuals with Noonan Syndrome note that treatments for cognitive and musculoskeletal abnormalities are a high priority (Bernardino et al., 2022; Doherty et al., 2023; Tiemens et al., 2023). Yet, it is difficult to discern whether functional changes are a result of aberrant signaling in neurons, myelinating oligodendrocytes, or complex interactions during development. Studies have shown a specific role for ERK1/2 signaling in regulating rodent myelinating glial development *in vivo* (Ishii et al., 2012; Ishii et al., 2013; Newbern et al., 2011), and pharmacological MEK1/2 inhibition rescues myelination deficits caused by RASopathic signaling in mature oligodendrocytes (López-Juárez et al., 2017; Mayes et al., 2013). Effects of upstream RASopathy mutations in SynGAP1, NF1, and RAS that activate multiple downstream cascades, selectively in neurons, have been explored (Clement et al., 2012; Cui et al., 2008; Kim et al., 2023; Rumbaugh et al., 2006). However, the neuron-autonomous effects of downstream ERK1/2 hyperactivation in developing cortical excitatory circuits are less clear. We previously reported subtype-specific functions for ERK1/2 in the development of cortical inhibitory and excitatory neurons with a series of Cre-dependent conditional mouse lines. Deletion of *Mek1/2* (*Map2k1/2*) in postmitotic excitatory neurons resulted in reduced corticospinal axon extension into the lumbar spinal cord followed by death of a subset of layer V CTIP2^+^ neurons in the neonatal motor cortex (Xing et al., 2016). In contrast, hyperactivation of ERK1/2 signaling in excitatory neurons via overexpression of a Cre-dependent, phosphomimic *MEK1^S217/221E^*construct did not alter CTIP2^+^ density, but led to reductions in corticospinal elongation by P3 (Xing et al., 2016).

Given the marked excitatory neuron subtype-specific outcomes associated with modifying ERK1/2 activity, we sought to understand whether MEK1 hyperactivation in excitatory neurons during development is sufficient to alter cortical axon outgrowth to multiple targets and ultimately affect cognition. We utilized a *Nex/Neurod6:Cre* mouse to conditionally express a hyperactive *MEK1^S217/221E^*construct in newborn, postmitotic cortical glutamatergic neurons, but not oligodendrocytes or cortical inhibitory neurons (Goebbels et al., 2006; Krenz et al., 2008). *MEK1^S217/221E^* selectively drives downstream hyperactivation of ERK1/2, independent of directly manipulating parallel RTK-linked cascades. Moreover, *Nex:Cre*-directed expression minimizes cellular contributions from glia and inhibitory neurons known to be modified by changes in ERK1/2 signaling (Angara et al., 2020; Holter et al., 2021; Kang and Lee, 2019; Knowles et al., 2023a; Knowles et al., 2023b). This approach will help identify the excitatory neuron-autonomous functions of selective hyperactivation of MEK-ERK1/2 signaling during long-range cortical circuit formation.

Our results show that conditional *MEK1^S217/221E^* overexpression leads to a persistent increase in phosphorylated ERK1/2 (pERK1/2) levels in axonal projections that coincides with reduced motor cortex derived axon arborization in intracortical and subcortical target domains. Moreover, mutant cortices exhibited reduced expression of a canonical immediate early gene, Arc, and deficits in learning a single-pellet retrieval task (Guo et al., 2015; Kawai et al., 2015; Peters et al., 2017; Whishaw et al., 1993). To test whether these deficits were cortical layer V-autonomous, we crossed the *MEK1^217/221E^* mouse line with a *Rbp4:Cre* mouse. Viral tracing analysis of contralateral corticocortical, corticostriatal, and corticospinal projections in *Rbp4:Cre MEK1^217/221E^* mutants revealed a significant decrease in axonal innervation, but no deficit in skilled motor learning. Collectively, these results indicate that excitatory neuron-specific hyperactivation of MEK1 in the cortex is sufficient to disrupt the development of cortical axonal projections, the expression of Arc, and select aspects of skilled motor learning.

## Materials and Methods

### Transgenic Mice

All animal experiments were performed in accordance with established procedures approved by the Institutional Animal Care and Use Committee of Arizona State University (Protocol# 22-1924R) and NIH guidelines for the use and care of laboratory animals. Mice used in this work were of a mixed genetic background, housed in standard conditions, and kept on a 12-hour light/dark cycle with *ad libitum* food and water. *NeuroD6/Nex-Cre (NeuroD6^tm1(cre)Kan^)* mice were kindly provided by Dr. Klaus Nave and Dr. Sandra Goebbels (MGI:2668659) (Goebbels et al., 2006). *Rbp4:Cre* mice were obtained from the MMRRC (RRID:MMRRC_031125-UCD) (Gong et al., 2007). *Mek1^S217/221E^ (Tg(CAG-cat,-Map2k1*)243Rbns)* mice were provided by Dr. Maike Krenz and Dr. Jeffrey Robbins (MGI:3822131) (Krenz et al., 2008). Cre-dependent *tdTomato/Ai9* (B6.-Gt(ROSA)26Sor^tom(CAG-tdTomato)Hze^/J) mice were ordered from Jackson Laboratories (RRID:IMSR_JAX:007909) (Madisen et al., 2010). Each experiment was replicated a minimum of three times from independent litters.

Genomic DNA was extracted from toe or tail samples and amplified with standard polymerase chain reaction techniques. Primers for gene amplification are as follows 5’-3’: Cre - GCTAAACATGCTTCATCGTCGG and GATCTCCGGTATTGAAACTCCAGC amplify a 645 base pair allele. *Mek1^S217/221E^* – GTACCAGCTCGGCGGAGACCAA and TTGATCACAGCAATGCTAACTTTC amplify a 600 base pair allele. *Ai9* – CTGTTCCTGTACGGCATGG and GGCATTAAAGCAGCGTATCC amplify a 196 base pair allele.

### Tissue Preparation

Transcardial perfusions were performed on mice using a 4% paraformaldehyde/PBS solution and dissected brains were post-fixed overnight at 4°C. Brains were then cryopreserved via serial incubations in 15% and 30% sucrose/PBS solutions prior to embedding in Tissue-Tek O.C.T. (Tissue-Tek 4583) and rapid freezing. Cryostat sections were collected in cold PBS for free-floating staining or mounted on Fisherbrand Superfrost Plus slides (Fisher Scientific 12-550-15) and air-dried prior to staining. For vibratome sectioned material, fixed adult brains were mounted in agarose and sections were collected in PBS prior to free-floating immunohistochemistry.

### Viral Injections and Image Quantification

P1 pups were removed from their home cage as a group and individually cryo-anesthetized on ice for 4-6 minutes. Viral injections were then performed immediately with 50nl of solution using a 32-gauge beveled needle fitted to a Hamilton 5µl neuros syringe mounted on a manipulator arm. After injections, pups were placed on a 37°C heated surface for recovery and returned to their home cage as a group. Adeno-Associated Viral vectors containing a Cre-dependent tdTomato construct under the control of a strong CAG promoter (AAV-CAG-FLEX-tdTomato, UNC Viral Vector Core) were used for viral tracing experiments. For quantification of virally labeled RFP^+^ axonal arborization in the forebrain, we first manually assessed the number of RFP^+^ neurons in M1 from confocal images of at least three sections near the center of the injection site. Confocal images of axonal target domains from at least three sections were automatically thresholded in ImageJ using the Otsu method (Otsu, 1979). The total number of RFP^+^ pixels within the target domain was normalized to the number of labeled M1 neurons within the injection site to provide a relative estimate of axonal arborization. To quantify axon elongation along the spinal cord, RFP^+^ immunolabeling in the cervical dorsal funiculus was normalized to immunolabeling in hindbrain corticobulbar tract (CBT). Representative images were cropped, adjusted for brightness and contrast, and inverted in Photoshop for clear visualization of axonal projections.

### Immunohistochemistry

Tissue sections were rinsed with PBS/0.1% Triton X-100 (PBST) and blocked with 5% normal donkey serum in PBST at room temperature for ∼1 hour. Primary antibodies were diluted in PBST/5%NDS and incubated overnight at 4°C with gentle agitation. Primary antibodies used were: rabbit anti-MEK1 (RRID:AB776273, Abcam ab32091), rabbit anti-phospho-p44/42 MAPK (ERK1/2) (Thr202/Tyr204) (RRID:AB_2315112, Cell Signaling Technologies 4370), rabbit anti-ERK2 (RRID:AB_732210, Abcam ab32081), rabbit anti-ARC (RRID:AB_887694, Synaptic Systems 156-003), mouse anti-β3Tubulin (TUJ1, RRID:AB_10063408, Bioloegend 801202), rabbit anti-RFP (RRID:AB_2209751, Rockland 600-401-379), chicken anti-RFP (RRID:AB_10704808, Rockland 600-901-379), and mouse anti-NeuN (RRID:AB_2298772, Millipore Sigma MAB377). After rinsing in PBS 3X, secondary antibodies diluted in PBST/5%NDS with DAPI (Sigma Aldrich 10236276001) were added and incubated overnight at 4°C with gentle agitation. Secondary antibodies include Alexa Fluor 488, 546, 568, and 647 conjugated to anti-rabbit IgG, anti-mouse IgG, or anti-chicken IgY (Invitrogen). Imaging was performed using a Zeiss LSM 800 or Leica SP5 laser scanning confocal microscope.

### Confocal Image Analysis and Quantification

The number of pERK1/2 labeled cells in M1 layer II/III were counted from 20x confocal images in at least two hemi-sections per mouse and averaged. We quantified the level of pERK1/2 immunolabeling in high resolution confocal images of layer V neurons collected with a 63X 1.4NA objective using the same acquisition settings. Individual NeuN^+^ pyramidal shaped somas >100 µm^2^ with a clear DAPI-labeled nucleus in cortical layer V of the motor cortex were randomly selected by a blinded observer. The integrated density of pERK1/2 immunolabeling in the cytoplasm and nucleus was then measured. A total of 68 control and 83 mutant neurons were analyzed from at least three sections per mouse, four mice per genotype, representing a total sampling area >14,000 µm^2^ per genotype.

To estimate the levels of pERK1/2 immunoreactivity in TUJ1^+^ callosal axons, standard confocal images of the corpus collosum were collected and analyzed. Standard confocal images of at least two tissue sections of the corpus collosum per mouse were collected and automatically thresholded in ImageJ in an unbiased manner using the default settings of Moments Autothreshold (Tsai, 1985). Colocalized pERK1/2^+^/TUJ1^+^ pixels were determined using Pierre Bourdoncle’s Colocalization Plugin with default settings and normalized to the total number of TUJ1^+^ pixels. Representative images were subsequently acquired in Airyscan mode using optimized settings to visualize the localization of pERK1/2 more clearly within TUJ1^+^ axons.

To quantify CST elongation in *RBP4:Cre; Ai9^+/-^* mice, confocal images of the dorsal funiculus in at least three lumbar segments per control and *MEK1^S217/221E^* expressing mice were autothresholded using Moments and the relative change in the number of RFP^+^ pixels was determined. Representative images have been cropped and adjusted for brightness and contrast in Photoshop for presentation.

### Western Blotting

Dorsal cortices were lysed in RIPA buffer (0.05M Tris-HCl, pH 7.4, 0.5M NaCl, 0.25% deoxycholic acid, 1% Triton X-100, and 1 mM EDTA, Millipore) and supplemented with 0.1% SDS, protease inhibitor (Sigma-Aldrich P2714), and phosphatase inhibitor cocktail II and III (Sigma-Aldrich P5726) via sonication. Lysates were then cleared by centrifugation and protein concentrations determined by Bradford Assay using a Peirce Protein Assay Kit (Thermo Scientific 23200). Equal amounts of protein were denatured in reducing sample buffer, separated by SDS-PAGE gels (BioRad 456-1035 and 456-8095), and transferred to PVDF membranes (BioRad 162-0177). Blots were blocked with 5% low-fat milk TBS containing 0.1% Tween-20 (TBST) for 1hr at room temperature, then incubated overnight at 4°C with primary antibodies in 4% BSA in TBST. Primary antibodies used were: rabbit anti-MEK1 (RRID:AB_776273, Abcam ab32091), rabbit anti-phospho-p44/42 MAPK (ERK1/2) (Thr202/Tyr204) (RRID:AB_2315112, Cell Signaling Technologies 4370), rabbit anti-ERK2 (RRID:AB_732210, Abcam ab32081), rabbit anti-ARC (RRID:AB_887694, Synaptic Systems 156-003), rabbit anti-GAPDH (RRID:AB_561053, Cell Signaling Technologies 2118), and rabbit anti-β-actin (RRID:AB_10950489, Cell Signaling Technologies 8457). After washing with TBST, membranes were incubated with donkey anti-rabbit HRP-conjugated secondary antibodies (Jackson Immunoresearch 711-035-152) in 5% milk/TBST for 2hr at room temperature. Membranes were washed with TBST and signal was detected with SuperSignal West Pico chemiluminescent (Thermo Scientific 32106) and exposed to radiographic film (Thermo Scientific 34089). Quantification of Western Blots was performed using a high-resolution scanned image of the exposed radiographic film in a grayscale TIFF file format. Bands were selected for quantification of integrated density in ImageJ following background subtraction from an unlabeled region and normalized to either the loading control or non-phosphorylated target.

### Open Field, Accelerating Rotarod, and Single-Pellet Retrieval

Open field testing was performed in an open top opaque plexiglass box measuring 38×38×30cm. The area within the box is illuminated by a centered, overhead 40-watt spotlight. Video recordings of open field behavior were scored by a blinded observer. Rotarod analysis was performed on a San Diego Instruments accelerating rotarod. Animals were tested for three trials per day for five consecutive days, accelerating the rotarod from 4 to 40 RPM with a 15min intertrial interval. The latency to fall was measured for each of three trials during a day and averaged.

A single-pellet retrieval task was performed using only male mice that were acclimated to sucrose pellets (TestDiet 1811555) in their home cages at least one week prior to testing. Mice were habituated for 10-15min per day, for two days in the testing room and chamber, which consists of a 15×10×20cm transparent acrylic box with a 5mm wide slit on the right and left side of one end that opens to a 10mm high food platform. During the subsequent two training days, mice were observed while single sucrose pellets were made regularly available during a 10-20min session. Only mice that attempted at least 20 reaches were further analyzed. Daily testing was performed for 10 min per day, for 5 days by placing a single pellet on the food platform. To enhance motivation for obtaining sucrose pellets, *Nex:Cre;MEK1^S217/221E^* mice were mildly food restricted by limiting free feeding to 2-6 hours daily while maintaining at least 90% of their initial weight. Due to unexpected negative health effects of mild food restriction in *RBP4:Cre; MEK1^S217/221E^* mice, these mutants and associated control mice were only restricted during a 2-4 hr period prior to a single 5 min session per day.

Attempts were scored as a “success” if the mouse was able to retrieve the sucrose pellet with its preferred paw and place it in its mouth. If the mouse grasped the pellet but failed to return it to its mouth, the attempt was scored as a “drop”. If the mouse clearly reached for the pellet but failed to grasp it, the attempt was categorized as a “miss”. For each mouse, the proportion of successful out of total attempts was calculated. Accelerating rotarod and single-pellet retrieval results were analyzed using a two-way repeated measures ANOVA in SPSS.

## Results

### Hyperactivation of MEK1 increases levels of pERK1/2 in cortical excitatory neuron axons

To investigate the developmental effects of excitatory neuron-autonomous hyperactive MEK-ERK1/2 signaling within the cortex *in vivo*, we generated mice expressing *Nex/NeuroD6:Cre*, Cre-dependent hyperactive *Mek1/Map2k1^S217/221E^*, and Cre-dependent tdTomato/RFP (*Ai9^+/-^*) (Goebbels et al., 2006; Krenz et al., 2008; Madisen et al., 2010). As expected, E14.5 *Nex:Cre; Ai9^+/-^* forebrains showed RFP expression in excitatory neurons and axons in the cortical plate (CP) and intermediate zone (IZ), but not in neural stem cells in the ventricular zone (VZ) (Fig. 1A-B) (Xing et al. 2016, Goebbels et al. 2006). Substantial MEK1 overexpression was apparent in recombined excitatory neuron somas and long-range axonal projections in *Nex:Cre; MEK1^S217/221E^; Ai9^+/-^* compared to controls (Fig. 1B-C, G-H). Western blots of dorsal cortical lysates from E14.5 *Nex:Cre; MEK1^S217/221E^* embryos demonstrate a 5.29 ± 0.93 fold-increase in MEK1 expression relative to controls (Mean ± SEM, n=5, Student’s t-test p=0.002) (Fig. 1K-L).

**Figure 1.**
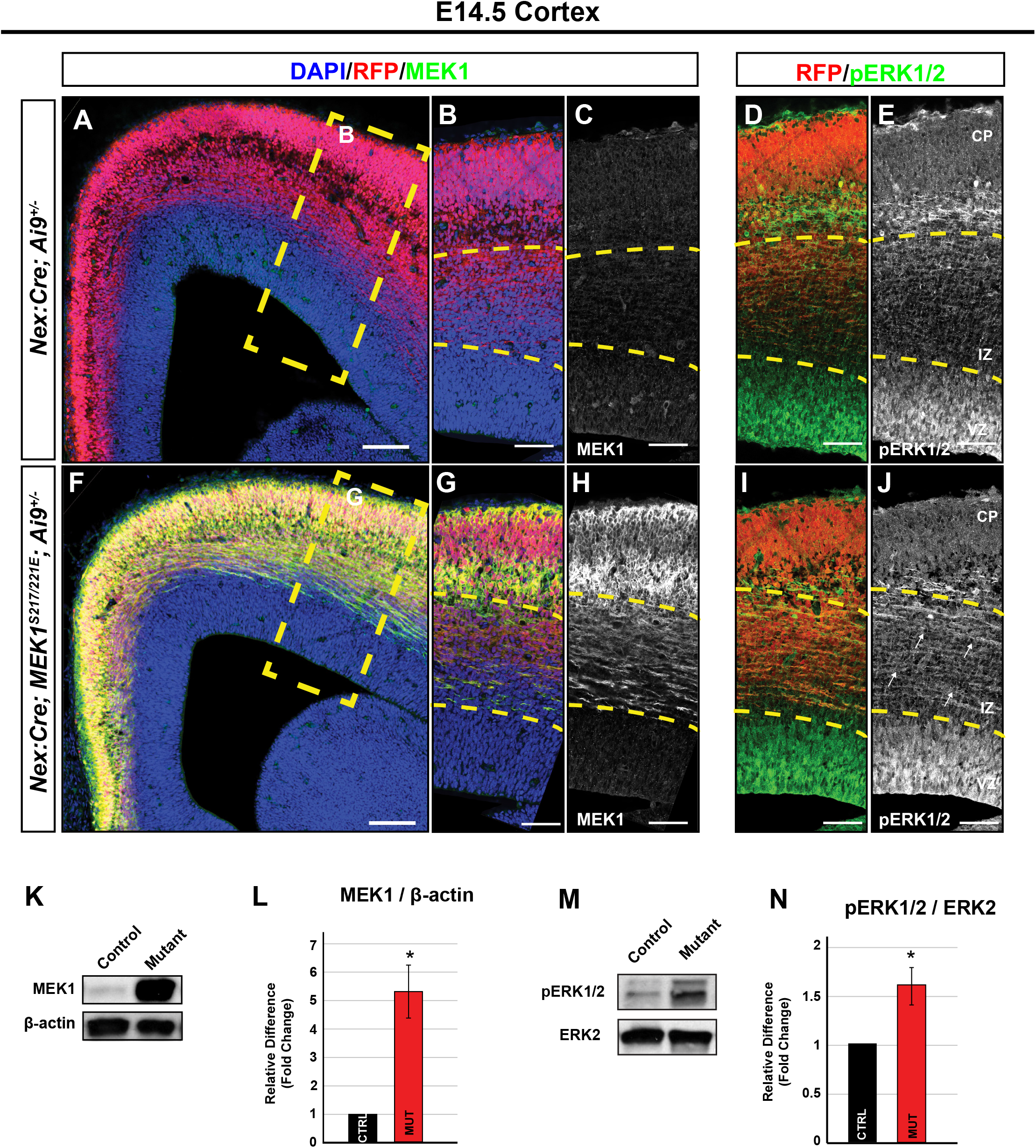
MEK1 hyperactivation selectively in immature glutamatergic neurons results in increased levels of pERK1/2 by E14.5. **A-J.** Immunolabeling of E14.5 *Nex:Cre; MEK1^S217/221E^; Ai9^+/-^* mutant cortices shows a substantial increase of MEK1 expression in the intermediate zone (IZ) and cortical plate (CP) of mutants (F-H) when compared to *Nex:Cre; Ai9^+/-^* controls (A-C) (n=4). RFP^+^ axons within the IZ of mutants (I-J, white arrow heads) exhibited a qualitative enrichment in levels of phosphorylated ERK1/2 relative to controls (D-E) (n=4). We did not detect a notable difference in the levels of pERK1/2 in the ventricular zone (VZ), where recombination is essentially absent in these mice. **K-N.** Western blots of E14.5 dorsal cortical lysates (K) reveal a 5-fold increase of MEK1 in mutants compared to controls (L) (Mean ± SEM, n=5, * = Student’s t-test p=0.002). We also observed a significant increase in pERK1/2 levels in mutants compared to controls (M-N) (Mean ± SEM, n=10, * = Student’s t-test p=0.005). Scale bars: A, F = 100 µm, B-E, G-J = 60µm.

*MEK1^S217/221E^* hyperphosphorylates ERK1/2 in cell free assays and in culture, but the extent of downstream effects *in vivo* varies in different tissue types (Alessi et al., 1994; Bueno, 2000; Cowley, 1994; Klesse et al., 1999; Krenz et al., 2008; Lajiness et al., 2014; Li et al., 2012). We directly measured whether excitatory neuronal *MEK1^S217/221E^* overexpression coincides with increased phosphorylated ERK1/2 (pERK1/2) levels *in vivo*. E14.5 dorsal cortical lysates from mutant mice exhibited a significant 1.61 ± 0.19 fold-increase in pERK1/2 levels compared to controls (Mean ± SEM, n=10, Student’s t-test p=0.005) (Fig. 1M-N). Immunolabeling for pERK1/2 in *Nex:Cre; MEK1^S217/221E^; Ai9^+/-^* mice revealed a relative qualitative enrichment of pERK1/2 in the intermediate zone and nascent RFP^+^ cortical excitatory neuron axons (Fig. 1I-J, white arrows), while comparable levels of pERK1/2 were observed in the unrecombined VZ of control and mutant embryos.

We next evaluated MEK1 and pERK1/2 activation in adult brains. Adult *Nex:Cre; MEK1^S217/221E^* cortical lysates revealed a statistically significant 1.61 ± 0.2 fold increase in MEK1 expression over controls (mean ± SEM, n=3, Student’s t-test p=0.041) (Fig. 2A-B). The proportion of excitatory neurons in whole adult cortical lysates is relatively less than at E14.5 and may contribute to the reduced fold-increase in MEK1 in adult mutants when compared to E14.5 mutants. We did not detect a significant difference in pERK1/2 levels between *Nex:Cre; MEK1^S217/221E^* and control whole cortical lysates (n=3, Student’s t-test p=0.832) (Fig. 2C-D). MEK1 immunolabeling of the cortex revealed a substantial increase in mutant excitatory neurons somas and axons within the corpus callosum when compared to the controls (Fig. 2E-H).

**Figure 2.**
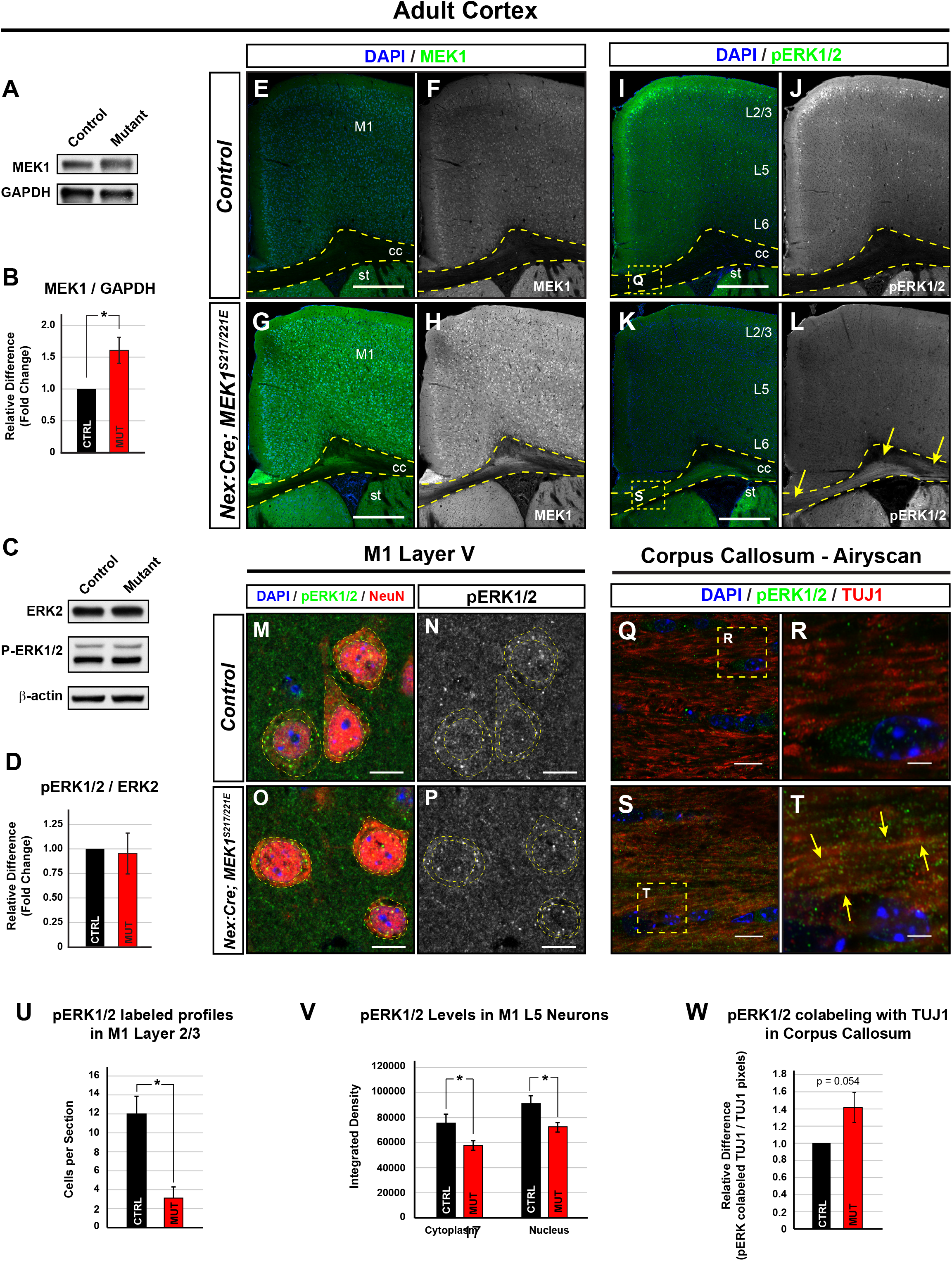
Increased levels of pERK1/2 in cortical axons of adult Nex:Cre; MEK1^S217/221E^ mice. **A-D.** Western Blots of adult cortical lysates revealed a 1.61 ± 0.20 fold increase in MEK1 in *Nex:Cre; MEK1^S217/221E^* mutants when compared to control mice (Mean ± SEM, n=3, Student’s t-test p=0.041) (A-B). No significant difference in pERK1/2 levels between mutant and control mice was detected in adult cortical lysates (Mean ± SEM, n=3, Student’s t-test p=0.389) (C-D). **E-W.** An increase of MEK1 immunolabeling was apparent in the cortex of adult *Nex:Cre; MEK1^S217/221E^* mice (G-H) relative to controls (E-F). Phosphorylated ERK1/2 immunolabeling was qualitatively increased in the corpus callosum (dotted yellow outline) in adult mutant mice (K-L, yellow arrows), when compared to controls (I-J). Within the grey matter, we noted a significant reduction in the number pERK1/2 labeled neurons in layer II/III of mutants (L, U) (Mean ± SEM, n=6, * = Student’s t test p=0.002). High resolution confocal imaging and analysis of NeuN^+^ pyramidal neurons in M1 layer V (M-P, dotted yellow outlines) revealed a decrease in cytoplasmic and nuclear levels of pERK1/2 levels (O-P, V) when compared to controls (M-N) (Mean ± SEM, n=68 control and 83 mutant neurons from four mice per group, * = Student’s t-test p<0.01). We detected a modest increase in pERK1/2/TUJ1 co-localization in the corpus callosum of mutants (S-T, yellow arrows) compared to controls (Q-R, quantified in W, Mean ± SEM, n=4, Student’s t test p = 0.054). Scale bars E,L = 500µm, M-P = 10µm, Q-T = 2 µm (M1 = primary motor cortex, cc = corpus callosum, st = striatum).

Control forebrains immunolabeled for pERK1/2 showed the expected enrichment in a subset of neuronal somas in layer II/III, with relative low levels in the corpus callosum (Fig. 2I-J) (Hart and Balleine, 2016; Lee et al., 2021). Interestingly, in *Nex:Cre; MEK1^S217/221E^* mutant mice, we observed a significant decrease in pERK1/2^+^ layer II/III neurons in M1 when compared to controls (n=6, Student’s t test p=0.002) (Fig. 2U). Moreover, semi-quantitative high-resolution analyses of pERK1/2 immunolabeling intensity in mutant layer V neurons revealed a reduction in the cytoplasm and the nucleus when compared to controls (n=68 control and 83 mutant neurons from four mice per genotype, Student’s t-test p<0.01) (Fig. 2M-P, V). In contrast, we noted a qualitative increase in pERK1/2-immunolabeling in a number of mutant forebrain white matter tracts, such as the corpus callosum (Fig. 2L, yellow arrows), fimbria, and stria terminalis (Fig. S1), which are enriched in axons derived from excitatory populations in the cortex, hippocampus, and amygdala, respectively. High resolution analysis of pERK1/2 co-localization with TUJ1, a marker of axons, in the corpus callosum, demonstrated the presence of pERK1/2 in axons and a modest increase in co-localization in mutant axons compared to controls (n=4, Student’s t test p = 0.054) (Fig. 2Q-T, W, yellow arrows). Collectively, these data suggest glutamatergic neuron-autonomous *MEK1^S217/221E^* expression results in significant hyperactivation of ERK1/2 in the axonal compartment of multiple forebrain projection subtypes.

### MEK1^S217/221E^ expression disrupts corticocortical and corticostriatal axon arborization

We previously reported a reduction in corticospinal tract (CST) elongation in *Nex:Cre; MEK1^S217/221E^* mice by postnatal day 3 (P3) that persisted to P30 (Xing et al., 2016). Here, we asked whether MEK1^S217/221E^ expression is sufficient to reduce axon arborization in two other target domains, the contralateral cortex and striatum. To unilaterally label developing neurons and their axons, a Cre-dependent tdTomato/RFP adeno-associated viral vector (AAV-*CAG-FLEX-tdTomato*) was injected into the primary motor cortex (M1) at P1 (Fig. 3A). There was no significant difference in the density of tdTomato-labeled cells per section between *Nex:Cre; MEK1^S217/221E^* (224.73 ± 26.73) and control (189.96 ± 14.20) motor cortices (mean ± SEM, n=5, Student’s t-test, p=0.284) (Fig. 3B-D). To estimate axonal arborization, tdTomato-labeled pixels in contralateral M1 and striatum were quantified. There was a statistically significant reduction of contralateral corticocortical arborization in *Nex:Cre; MEK1^S217/221E^*mice (631.68 ± 27.51) compared to controls (791.79 ± 42.88) when normalized to number of transduced cells in M1 (Mean ± SEM, n=5, Student’s t-test, p=0.014) (Fig. 3E, F, H, I, K). Contralateral corticostriatal projections into the dorsolateral striatum were also significantly reduced in the *Nex:Cre;MEK1^S217/221E^* mice (control: 255.67 ± 23.64, mutant: 155.44 ± 21.18, Mean ± SEM, n=5, Student’s t-test, p=0.013) (Fig. 3E, G, H, J, L). Together with our previous findings, cortical glutamatergic neuron autonomous *MEK1^S217/221E^* expression appears sufficient to hyperactivate ERK1/2 during axonal development and reduce arborization into multiple long-range targets.

**Figure 3.**
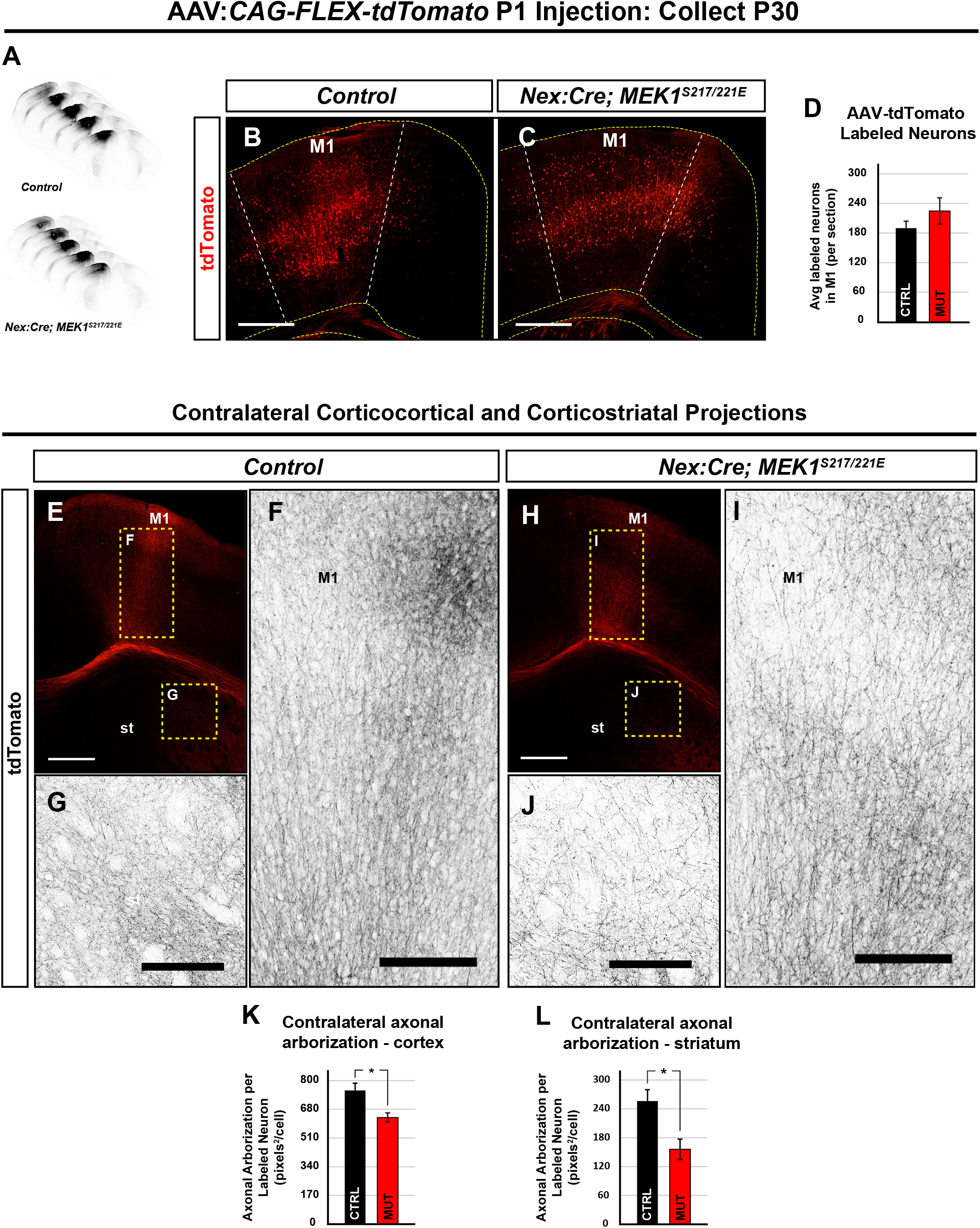
Nex:Cre; MEK1^S217/221E^ mice exhibit disrupted contralateral corticocortical and corticostriatal arborization. **A-D.** An adeno-associated viral vector, AAV:*CAG-Flex-tdTomato* was injected into the left primary motor cortex at P1 and brains were collected at P30 as shown in the rostral to caudal serial sections surrounding the injection site (A) and representative images of the injected region in the primary motor cortex (M1) of the *Nex:Cre; MEK1^S217/221E^* (C) and control (B) forebrains. No significant difference in the average density of RFP-labeled cells was detected between *Nex:Cre; MEK1^S217/221E^*and control mice (Mean ± SEM, n=5, Student’s t-test p=0.2839) (D). **E-L.** Representative confocal images of contralateral corticocortical and corticostriatal (st = striatum) RFP-labeled axonal arborization in the *Nex:Cre; MEK1^S217/221E^* (H-J) and control mice (E-G). *Nex:Cre; MEK1^S217/221E^* mice showed a 19.57% ± 4.30 reduction of normalized intracortical axon arborization (I) compared to control mice (Mean ± SEM, n=5, Student’s t-test p=0.014) (F, quantified in K). Cortical axon arborization in the mutant dorsolateral striatum was reduced 37.50% ± 9.14 (J) compared to control mice (Mean ± SEM, n=5, Student’s t-test p=0.013) (G, quantified in L). Scale bars: B, C, E, H = 500µm, F, G, I, J= 100µm.

### ARC protein levels are reduced by MEK1 hyperactivation

ERK1/2 is a known regulator of glutamatergic signaling and activity dependent plasticity (Bramham et al., 2010; Thomas and Huganir, 2004). Whether *MEK1^S217/221E^* expression alters circuit-level cortical activity in adults is unclear. Activity-Regulated Cytoskeleton-associated gene (ARC) is an immediate early gene (IEG) triggered by glutamate receptor signaling via ERK1/2 and other pathways (Cao et al., 2015; Chotiner et al., 2010; Correa et al., 2012; Ebert and Greenberg, 2013; Pintchovski et al., 2009; Ying et al., 2002). Our data shows axonal arborization, somal and nuclear levels of pERK1/2, are reduced in *Nex:Cre; MEK1^S217/221E^* mutants, suggesting that glutamatergic drive may be diminished. Indeed, western blots of adult whole cortical lysates revealed a significant 37 +/-11% decrease in ARC expression in *Nex:Cre;MEK1^S217/221E^* mice compared to controls (n=11, Student’s t-test p<0.005) (Fig. 4A-B). Immunolabeling further indicated a widespread decrease in cortical ARC levels *in vivo* (n=4) (Fig. 4C-F). Together, these findings support that developmental expression of hyperactive MEK1 in glutamatergic neurons reduces cortical ARC expression *in vivo*.

**Figure 4.**
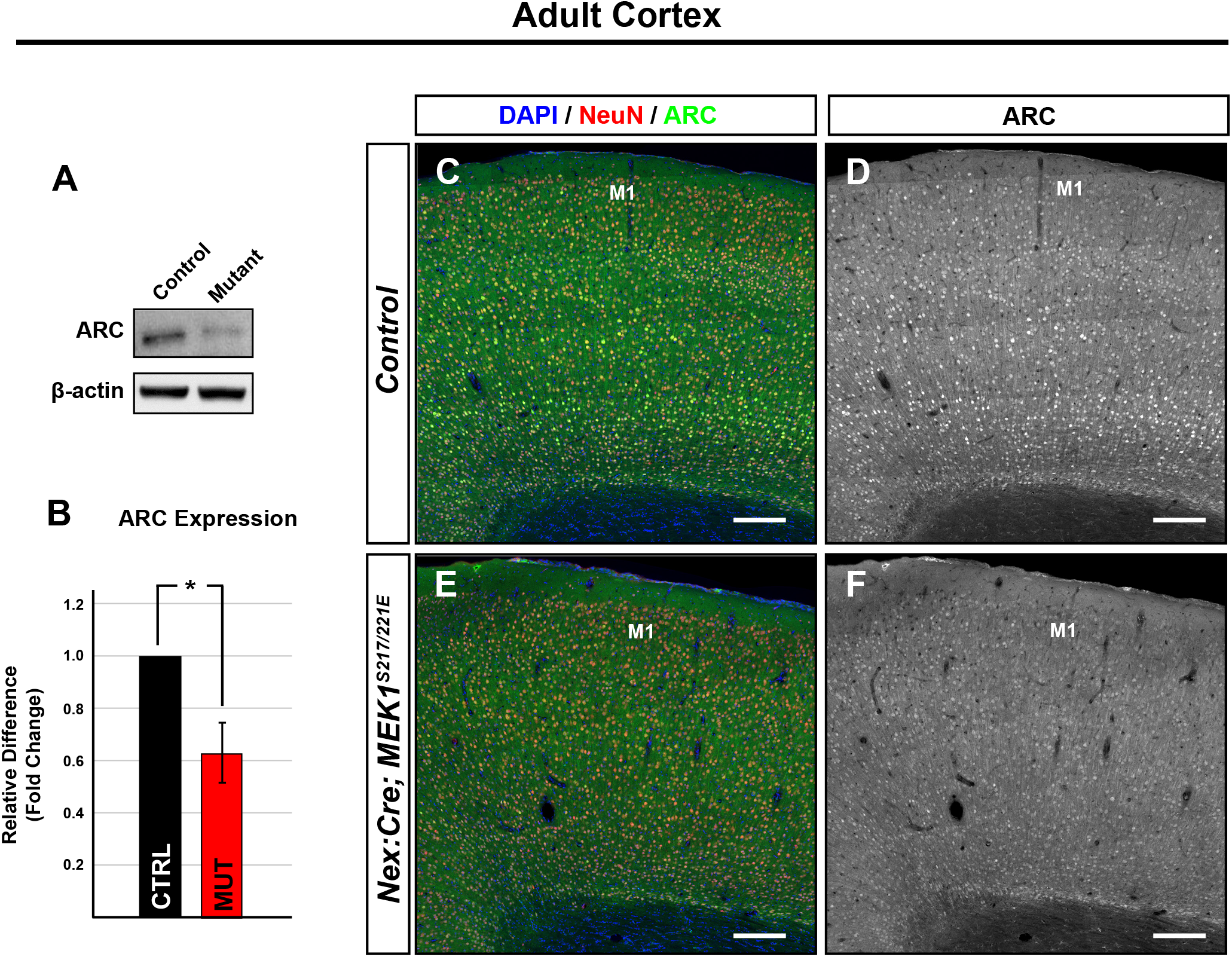
MEK1^S217/221E^ reduces cortical ARC protein expression. **A-B.** Western Blots of adult cortical lysates showed a significant reduction in ARC in *Nex:Cre; MEK1^S217/221E^* mutants when compared to control mice (Mean ± SEM, n=11, Student’s t-test, p<0.005). **C-F.** Immunolabeling revealed a qualitative reduction of ARC in *Nex:Cre; MEK1^S217/221E^* cortices (E-F) when compared to controls (C-D, n=5). Scale bars = 200um.

### Motor skill acquisition is disrupted in Nex:Cre; MEK1^S217/221E^ mice

Common phenotypes in RASopathic individuals include hypotonia, changes in motor coordination and altered motor learning (Blüthgen et al., 2017; Davis et al., 2000; Monje et al., 2005). We thus asked if hyperactivation of MEK1 in *Nex:Cre* expressing cells was sufficient to drive changes in adult mouse motor behavior in open field, accelerating rotarod, and single-pellet retrieval assays (Crawley, 1999). Open field testing revealed no significant difference in total distance traveled between *Nex:Cre; MEK1^S217/221E^* and control mice (Fig. 5A, n=12, Student’s t-test, p=0.075). Time spent in the center of the open field arena is a gauge of anxiety-related behavior, however, we did not observe a significant difference (Fig. 5A, n=12, Student’s t-test p=0.21). We also did not observe a significant effect of genotype during accelerating rotarod testing, which assesses motor coordination and balance (Fig. 5B, [F(1, 27)=0.246, p=0.624], n=15 controls & 14 mutants). These data reveal that *Nex;Cre; MEK1^S217/221E^* mice have relatively normal global voluntary locomotor behavior, motor coordination, and balance.

**Figure 5.**
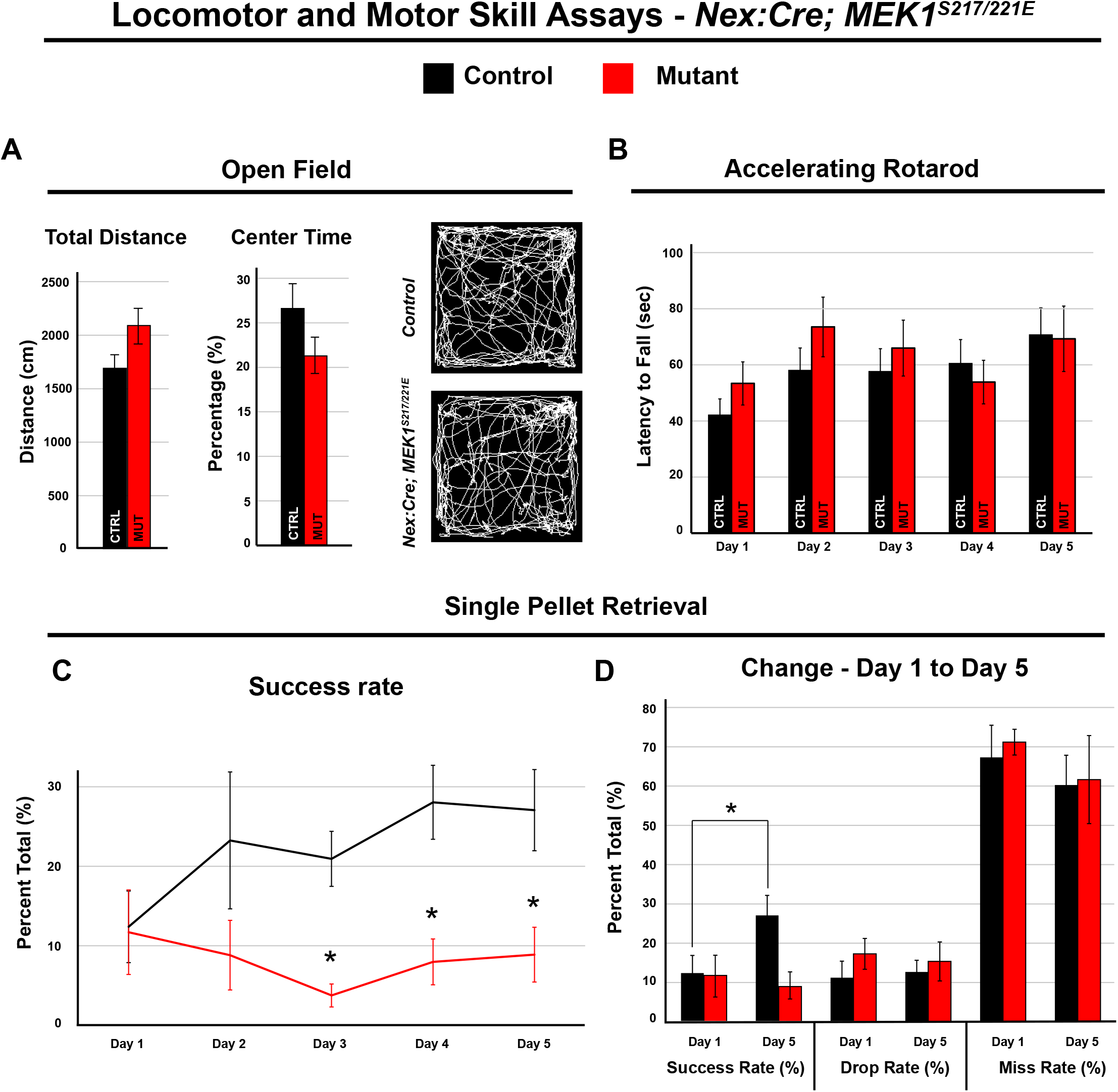
Motor skill acquisition is disrupted by MEK1 hyperactivation in cortical glutamatergic neurons. **A.** Open field testing of *Nex:Cre; MEK1^S217/221E^*(n=12) and control (n=12) mice revealed no significant difference in distance traveled (Mean ± SEM, Student’s t-test p=0.075) or time spent in the center of the arena (Mean ± SEM, Student’s t-test p=0.21) (A). **B.** Accelerating rotarod testing revealed a significant effect of day [F(4, 27)=6.144, p<0.001] over 5 days of testing, but no significant effect of genotype [F(1, 27)=0.246, p=0.624] (Mean ± SEM, n=15 controls & 14 mutants) (B). **C-D.** In the single-pellet retrieval task, the control success rate significantly improved over five days from 12.39 ± 4.50 to 27.06 ± 5.11 percent success rate (Mean ± SEM, Students t-test D1 vs D5 p=0.04, n=11) (C-D). The *Nex:Cre; MEK1^S217/221E^* mutant mice exhibited a significantly lower success rate relative to controls during the final three days of testing [F(1,16)=11.04, p=0.004] (C-D, Mean ± SEM, n=11 controls & 8 mutants, * = LSD post-hoc p < 0.05).

We next utilized a single-pellet retrieval task to determine if there was a deficit in fine forelimb motor function and learning, which relies heavily on the motor cortex (Chen et al., 2014; Kawai et al., 2015; Li and Hollis, 2021). This task assays the ability of mice to reach through a narrow slot with a single forelimb to retrieve a sucrose pellet. We measured the percentage of successful retrievals over the total number of reaching attempts across 5 consecutive days of testing. Importantly, we found that *Nex:Cre; MEK1^S217/221E^* mutants exhibit a significant decrease in success rate relative to controls (Fig. 5C, main effect of genotype [F(1,16)=11.04, p=0.004], n=11 controls & 8 mutants). The success rate in mutant mice was significantly decreased relative to controls during the last three days of testing (* = LSD post-hoc p < 0.05). However, no significant difference in success rate was noted on the first day of testing (LSD post-hoc p=0.59) or in the number of attempts [F(1,16)=0.06, p=0.81]. Collectively, these results indicate that the *Nex:Cre; MEK1^S217/221E^* mice do not exhibit initial, basal differences in fine forelimb motor function, but display a significant impairment in learning a skilled forelimb reaching and grasping task.

### Mouse model for studying layer V autonomous effects of ERK1/2 signaling

Since *Nex:Cre* recombines in all cortical layers, it is unclear whether axonal deficits are due to direct layer V specific effects or changes in upstream layers in the cortical circuit. To determine the layer V autonomous functions of ERK1/2 signaling, we employed a *Rbp4:Cre* mouse that drives recombination primarily in cortical layer V neurons in the cortex (Gong et al., 2007; Kozorovitskiy et al., 2012) and outside of the nervous system in the liver (Steinhoff et al., 2021; Thompson et al., 2017). Inducible deletion of *Erk1/2* in adult mice has been shown to trigger hepatocyte death and increased mortality, while ERK1/2 hyperactivation induces hepatocyte proliferation and carcinoma (Cingolani et al., 2022; Huynh et al., 2003; Ito et al., 1998). Attempts to generate *Rbp4:Cre; Erk1^-/-^; Erk2^fl/fl^* loss-of-function mice did not result in viable births (0 of 85, expected 1:4 births). However, *Rbp4:Cre; MEK1^S217/221E^; Ai9^+/-^* mice are viable into adulthood, but exhibit hepatomegaly (Fig. S2M). Nonetheless, *Rbp4:Cre; MEK1^S217/221E^; Ai9^+/-^* mice express RFP and elevated levels of MEK1 in cortical layer V somas and axon projections and provide a model for evaluating layer V-autonomous effects (Fig. S2D-F, J-L).

### Layer V directed expression of MEK1^S217/221E^ reduces cortical axon arborization

We did not observe a decrease in RFP^+^ cortical layer V neuron density in M1 of *Rbp4:Cre; MEK1^S217/221E^; Ai9^+/-^* mutants relative to controls (Control = 372 ± 9, Mutant = 477 ± 38, Mean ± SEM, Student’s t-test p=0.054), consistent with past findings assessing the proportion of layer V CTIP2^+^ neurons in *Nex:Cre; MEK1^S217/221E^* mice (Xing 2016) (Fig. S2B,E). We next asked if hyperactive MEK1 expression in layer V is sufficient to reduce axon extension. All layer V derived subcortical projections in the spinal cord dorsal funiculus were labeled by the Cre-dependent tdTomato/RFP reporter, *Ai9^+/-^*, in *Rbp4:Cre* mice. We observed that *Rbp4:Cre; MEK1^S217/221E^; Ai9^+/-^* mice had a 28 +/-8% reduction in RFP^+^ pixel density in the lumbar dorsal funiculus compared to controls (Fig. 6A-G) (Mean ± SEM, n=5, Student’s t-test p < 0.05).

**Figure 6.**
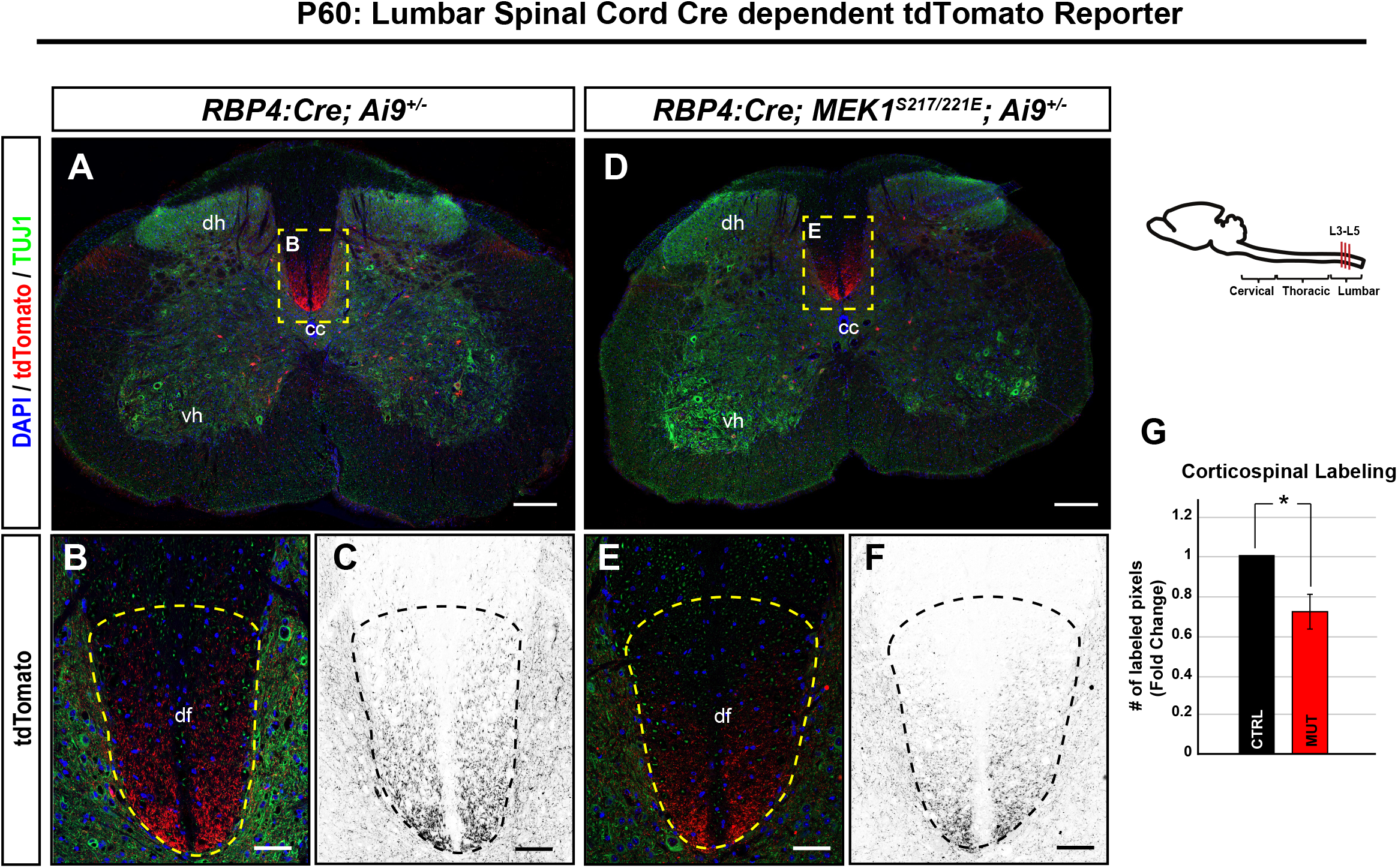
MEK1^S214/221E^ expression in cortical layer V reduces corticospinal axon elongation in the spinal cord dorsal fasciculus. **A-G.** *RBP4:Cre; Ai9^+/-^* control (A) and *RBP4:Cre; MEK1^S17/221E^; Ai9^+/-^* mutant (D) mice label descending cortical layer V derived corticospinal axons (dotted yellow box). Cross-sectional lumbar segments reveal a statistically significant 28% reduction of RFP^+^ labeled pixels in the dorsal funiculus of *RBP4:Cre; MEK1^S217/221E^; Ai9^+/-^* mice (E-F, dotted yellow outline) compared to *RBP4:Cre; Ai9^+/-^* controls (B-C, quantification in G, mean ± SEM, n=5, * = Student’s t-test p < 0.05). Scale bars A, D = 200um, B-C, E-F = 20um (dh = dorsal horn, cc = central canal, vh = ventral horn, df = dorsal funiculus).

The widespread bilateral labeling of all cortical layer V neurons in *Rbp4:Cre; Ai9^+/-^* mice made it challenging to determine the anatomical source of axonal labeling. Therefore, we utilized focal neonatal injection of a Cre-dependent tdTomato/RFP AAV into M1 to unilaterally label neurons at birth. Comparable numbers of neurons were labeled in M1 in control and *Rbp4:Cre; MEK1^S217/221E^* mice at P30 (Mean ± SEM, n=5, Student’s t-test p=0.400) (Fig. S3C). Although there was no difference in RFP labeling in corticobulbar tract (CBT), mutants had reduced RFP labeling in the cervical dorsal funiculus compared to controls, consistent with decreased CST elongation (Fig. 7A-E, Fig. S3D-H). Intracortical arborization in contralateral M1 of *Rbp4:Cre; MEK1^S217/221E^* mice was also significantly reduced when compared to control mice (Mean ± SEM, n=5, Student’s t-test p=0.019) (Fig. 7F-G, J-K, quantification in N). Contralateral dorsolateral striatal arborization was also significantly reduced in *Rbp4:Cre;MEK1^S217/221E^* mice when compared to controls (Mean ± SEM, n=5, Student’s t-test p=0.007) (Fig. 7H-I, L-M, quantification in O). Collectively, these results indicate that hyperactive ERK1/2 signaling in cortical glutamatergic layer V neurons is sufficient to reduce axonal elongation of multiple projection neurons.

**Figure 7.**
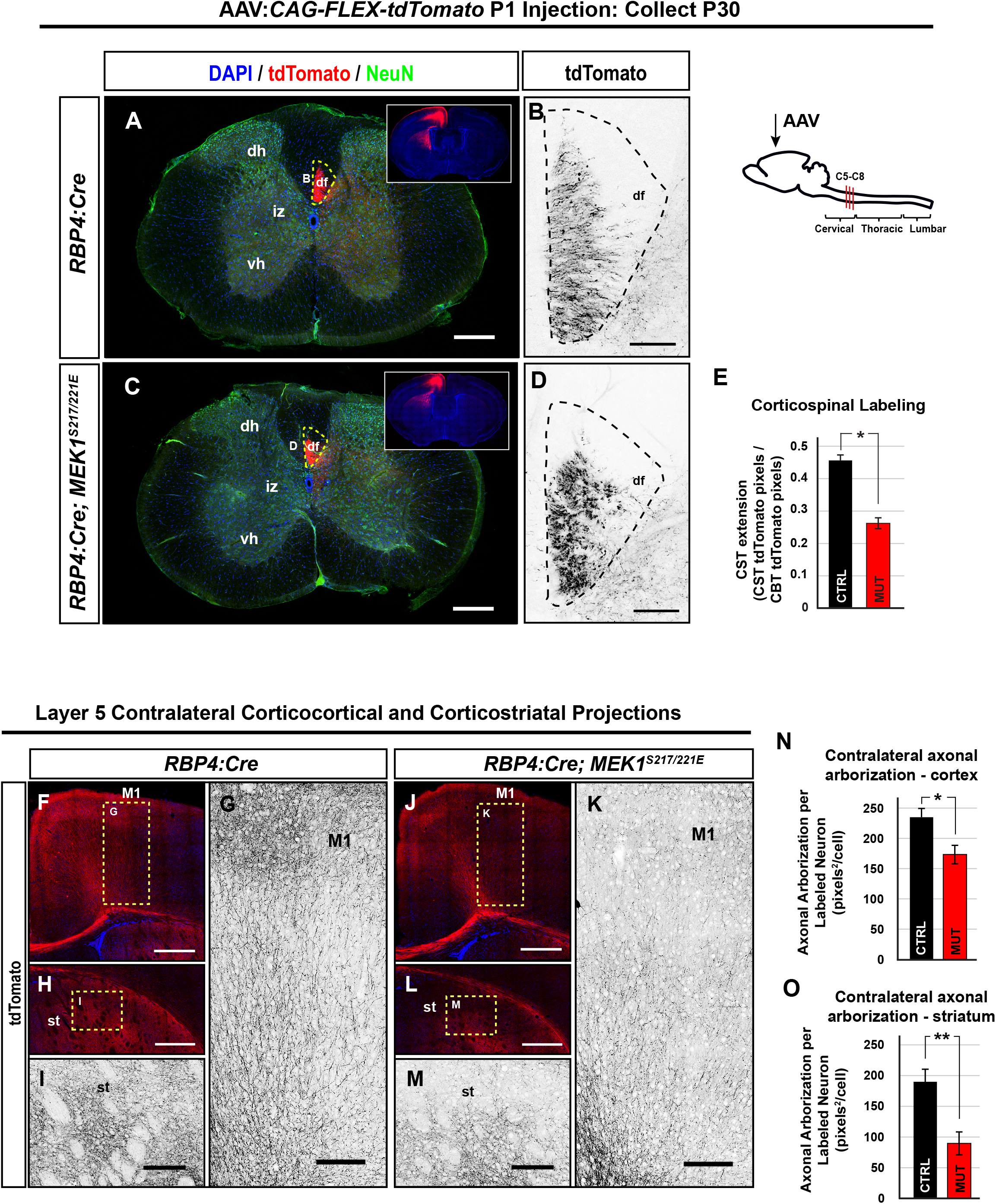
Corticospinal, corticocortical and corticostriatal layer V projections are reduced in *RBP4:Cre; MEK1^S217/221E^*mutants. **A-E.** Unilateral AAV labeling of descending corticospinal axons derived from M1 in RBP4:Cre mice. Cross sections of cervical spinal cord segments reveal a reduction of descending RFP^+^ corticospinal axons in the dorsal funiculus of *RBP4Cre; MEK1^S217/221E^* mutants (D) compared to *RBP4:Cre* control mice (B) when normalized to corticobulbar tract labeling (E, Mean ± SEM, n=3, Student’s t-test p=0.036). **F-O.** Virally-labeled *RBP4Cre; MEK1^S217/221E^* axons had significantly reduced arborization in the contralateral motor cortex (K) compared to *RBP4:Cre* controls (G) when normalized to the number of transduced cells in M1 (quantification in N, Mean ± SEM, n=5, * = Student’s t-test p=0.019). Additionally, *RBP4Cre; MEK1^S217/221E^* mice had significantly reduced arborization of the dorsolateral striatum (M) compared to *RBP4:Cre* controls (I) (quantification in O, Mean ± SEM, n=5, * Student’s t-test p=0.007). Scale bars: A,C, =200µm, B,D,= 20µm, F,H,J,L = 500µm, G,I,K,M = 150µm (dh = dorsal horn, df = dorsal funiculus, iz = intermediate zone, vh = ventral horn, M1 = primary motor cortex, st = striatum).

### Reduction in cortical wiring of layer V is expendable for skilled motor learning

Since *Rbp4:Cre;MEK1^S217/221E^* mice exhibit reduced axonal extension of cortical projection neurons, we tested whether these mutants display deficits in motor behavior. Hepatomegaly was observed in *Rbp4:Cre;MEK1^S217/221E^*mice and mouse models of chronic liver disease have been shown to exhibit hypolocomotion in the open-field, but little change in motor learning or anxiety-like phenotypes (Jover et al., 2006). When compared to control mice, *Rbp4:Cre;MEK1^S217/221E^*mutants showed a significant decrease in voluntary distance traveled in the open-field assay (n=22 controls and 11 mutants, Student’s t-test p=0.015) and a decrease in the proportion of time spent in the center (Student’s t-test p=0.001) (Fig. 8A). In contrast, no significant effect of genotype was detected on the accelerating rotarod task ([F(1,36)=0.007, p=0.933], n=26 controls & 12 mutants) (Fig. 8B).

**Figure 8.**
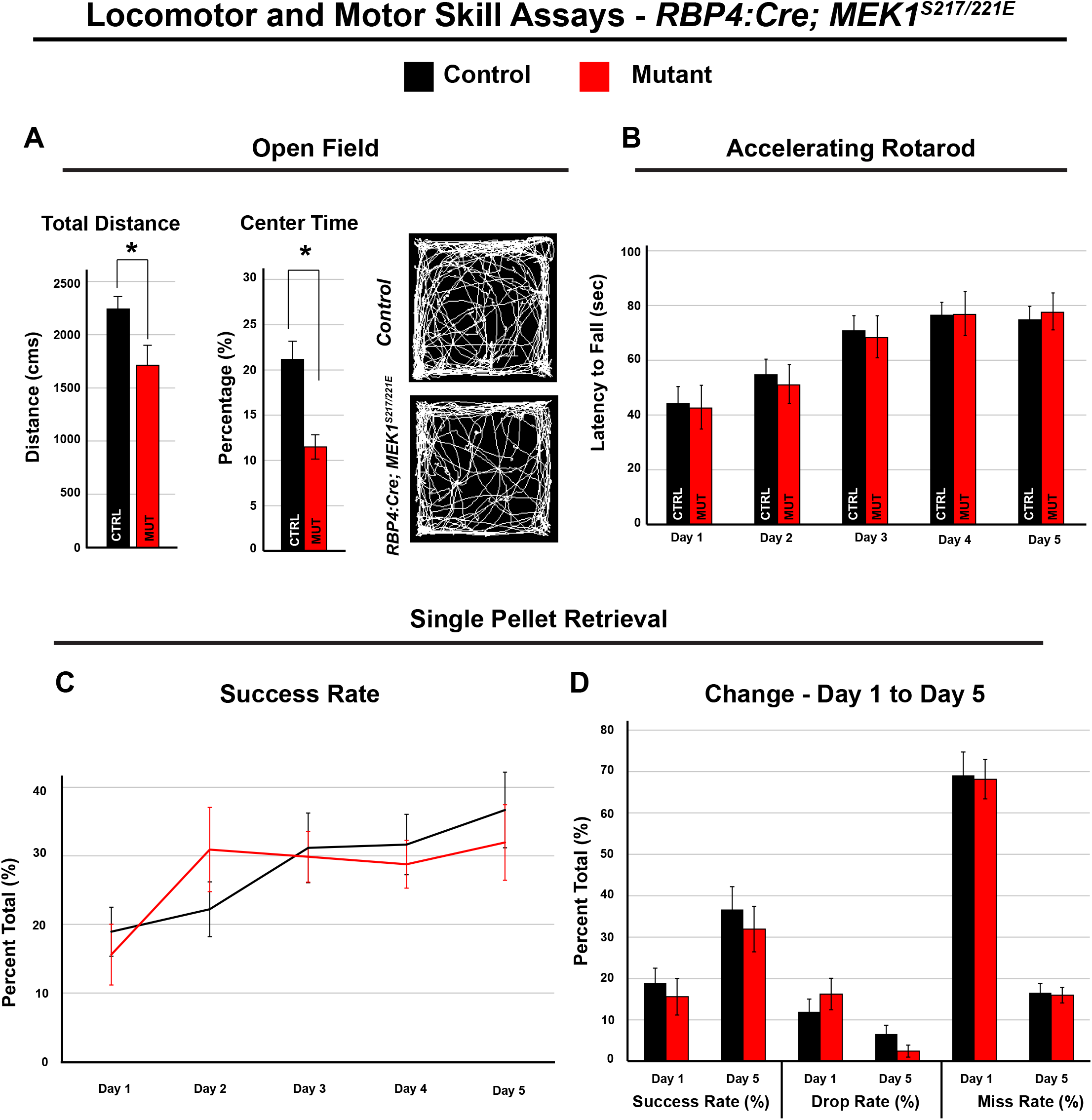
MEK1^S217/221E^ in cortical layer V glutamatergic neurons does not disrupt motor learning. **A.** Open field testing of RBP4*:Cre; MEK1^S217/221E^* and control mice revealed that mutants exhibit a significant reduction in total distance traveled (Mean ± SEM, n= 22 controls & 11 mutants, Student’s t-test p=0.01) and time spent in center of box (Mean ± SEM, n= 22 controls & 11 mutants, Student’s t-test p=0.001). **B.** Accelerating rotarod testing revealed no significant effect of genotype over 5 days of testing control and mutant mice ([F(1,36)=0.007, p=0.933] (Mean ± SEM, n=26 controls & 12 mutants). **C-D.** In the single-pellet retrieval task, there was no significant effect of genotype on daily success rate [F(1,24)=0.03, p=0.88] (C, Mean ± SEM, n=14 controls & 12 mutants). Both mutants and controls improved their success rate over five days [F(4,24)=5.81, p<0.01] (D, Mean ± SEM, n = 14 controls & 12 mutants).

In the single-pellet retrieval task, we found that mild food restriction in *Rbp4:Cre; MEK1^S217/221E^* mice resulted in health issues of unknown origin and spontaneous lethality. Thus, we substantially reduced the period of food restriction relative to our *Nex:Cre* experiments and only analyzed reaching and grasping behaviors during the first five minutes of each trial when mice were highly motivated. In stark contrast to *Nex:Cre; MEK1^S217/221E^* mutants, we found no significant effect of genotype on the success rate between control and *Rbp4:Cre;MEK1^S217/221E^*mice (Fig. 8C-D, ([F(1,24)=0.03, p=0.88], n=14 controls & 12 mutants). Taken together, these results indicate that layer V specific hyperactivation of MEK1 in glutamatergic neurons is insufficient to disrupt motor skilled learning.

## Discussion

Resolving the cell-specific effects of pathological ERK1/2 activity may help inform the rational design of therapeutic approaches for multiple neurodevelopmental conditions. Here, we tested whether cortical excitatory neuron-directed expression of hyperactive MEK1^S217/221E^ during early development modulates long-range axon outgrowth from sensorimotor cortices, expression of ARC, and motor-related behaviors. MEK1 hyperactivation selectively mimics a core downstream feature of multiple RASopathies without direct modulation of multiple RTK-linked signaling cascades associated with upstream mutations. We generated mice expressing Cre-dependent MEK1^S217/221E^ and *Nex:Cre* to target cortical excitatory neurons, but not cortical GABAergic neurons or glia. Axonal levels of pERK1/2 are persistently increased in these mutants. Importantly, ERK1/2 hyperactivation coincides with reductions in corticocortical, corticostriatal, and corticospinal axon projections arising from motor cortices. We also observe lower expression of the activity-dependent gene, Arc, within the mature cortex of mutant mice. While many basic motor behaviors remain intact, skilled motor learning in a reaching task is inhibited in these mutants. Lastly, we demonstrate that a layer V-specific MEK1^S217/221E^ mutant mouse recapitulates axonal outgrowth deficits, but motor learning is not altered. Overall, our data show that layer V autonomous expression of MEK1^S217/221E^ disrupts the development of long-range cortical axon projections, while MEK1 hyperactivation in developing *Nex:Cre*-expressing neurons appears necessary for diminished skilled motor learning (Fig. S4).

### Compartment-specific ERK1/2 activation in *MEK1^S217/221E^*-expressing neurons

*In vitro* studies have been vital for quantitative studies of kinase signaling and pharmacological inhibitors, but the precise effect of RASopathy mutations on pERK1/2 levels and distribution are more challenging to measure *in vivo*. We generated mutants with increased hyperactive MEK1 levels in developing excitatory neurons and observed increased levels of pERK1/2 in axonal, but not somal and nuclear compartments. Recent studies have shown that callosal layer II/III projections have enriched ERK2/MAPK1 protein within axonal growth cones (Engmann et al., 2022; Poulopoulos et al., 2019; Shigeoka et al., 2016). It will be important to explore whether enrichment of ERK1/2 pathway components within the axon renders this compartment relatively vulnerable to RASopathic mutations. High resolution measurement of RASopathic kinase activation in distinct subcellular domains *in vivo* may further illuminate core pathological mechanisms.

Paradoxically, the somal and nuclear levels of pERK1/2 were reduced in adult mutant excitatory neurons. A similar phenomenon has been observed in Drosophila MEK1 mutants, possibly due to early changes in the expression of morphogens during development (Goyal et al., 2017). The compensatory mechanisms that appear to override the biochemical hyperactivity of the MEK1^S217/221E^ mutant in neuronal soma and nuclei may involve ERK1/2-directed phosphatases enriched within somal compartments or developmental changes in upstream ERK1/2 activators (Caunt et al., 2008; Jeffrey et al., 2007; Li et al., 2007; Lim et al., 2015; Parmar et al., 2015; Poulopoulos et al., 2019). Nevertheless, pharmacological inhibitors of the ERK1/2 signaling cascade are actively being explored for clinical use in RASopathies (Guilding et al., 2007; Hatzivassiliou et al., 2010; Morris et al., 2013; Nakamura et al., 2009; Zamorano et al., 2018). For example, MEK1/2 inhibitors during postnatal stages reverses cardiac deficits and tumorigenesis in RASopathy models and patients with Noonan Syndrome (Andelfinger et al., 2019; Hilal et al., 2023; Wu et al., 2011). However, if developmental compensatory mechanisms lead to paradoxically reduced levels of pERK1/2 in select neuronal compartments, it is unclear whether pharmacological inhibition would be beneficial. Additional quantitative analyses of compartment-specific changes in pERK1/2 levels in RASopathy models may help inform the use of systemic pharmacological inhibitors.

### MEK-ERK1/2 hyperactivity reduces cortical long-range axon arborization

MRI-based studies of individuals with NF1 and Noonan Syndrome have discovered abnormalities in both functional and structural connectivity (Fattah et al., 2021; Filippi et al., 2013; Rai et al., 2023). While oligodendrocyte mediated changes in the myelination of axons are likely involved, the neuron-autonomous contributions to aberrant connectivity are less understood. We show that neuron-specific MEK1^S217/221E^ expressing mice did not exhibit significant loss of cortical layer V neurons, but layer V-autonomous hyperactivation of ERK1/2 reduces the arborization of corticocortical, corticostriatal, and corticospinal axons. *In vitro* studies have demonstrated ERK1/2 is necessary for aspects of trophic factor-induced axon outgrowth (Markus et al., 2002; Ozdinler and Macklis, 2006), axon guidance (Bagnard et al., 2004; Forcet et al., 2002; Yun et al., 2005), cytoskeletal organization (Atwal et al., 2003; Goold and Gordon-Weeks, 2005; Ray and Sturgill, 1987) and local protein translation in growth cones (Campbell and Holt, 2003). Whether hyperactive ERK1/2 activity disrupts these molecular processes or modulates other intracellular events associated with axon outgrowth *in vivo* is unclear. Axonal outgrowth in humans occurs during gestational stages, which makes therapeutic intervention more difficult. RASopathies are often diagnosed within the first years of life (Zenker et al., 2022). Therefore, it may be beneficial to study postnatal stages of axonal refinement and pruning in RASopathies, which may be more accessible to current treatment strategies.

Animal models of upstream RASopathic mutations in NF1, Ptpn11, and Ras have observed changes in multiple RTK-linked downstream intracellular cascades, including PI3K/Akt, JAK/STAT, PKC, mTOR and ERK1/2, and inhibitors of multiple cascades are capable of reversing certain cellular deficits (Anastasaki and Gutmann, 2014; Brown et al., 2012; Langdon et al., 2012; López-Juárez et al., 2017; Titus et al., 2017). Our work now shows that downstream MEK-ERK1/2 hyperactivity is sufficient to reduce long range axon arborization *in vivo*. Understanding whether upstream mutations also drive neuron-autonomous deficits in long-range excitatory neuron axonal outgrowth and may help inform the design of personalized, mutation-specific approaches to RASopathy therapeutics.

### Nex:Cre; MEK1^S217/221E^ mice exhibit decreased Arc and deficits in skilled motor learning

Activity-regulated cytoskeletal gene *Arc* is an immediate early gene that rapidly responds to neuronal activity and regulates synaptic plasticity (Steward and Worley, 2001; Steward et al., 1998). *Arc* has been shown to be dysregulated in Fragile X and Angelman Syndrome (Greer et al., 2010; Zalfa et al., 2003), however, whether RASopathic individuals exhibit aberrant *Arc* expression is not clear. Several reports show that glutamate and BDNF drive Arc transcription and translation via ERK1/2 (Chotiner et al., 2010; Correa et al., 2012; Nikolaienko et al., 2017; Pintchovski et al., 2009; Rao et al., 2006; Ying et al., 2002). Notably, our results indicate that MEK1 hyperactivation during early neuronal development unpredictably resulted in reduced phosphorylated ERK1/2 levels in the soma and nucleus of adult neurons. It is perhaps not surprising that ARC protein is also decreased in adult mutant cortices, in agreement with our past report showing P21 *Nex:Cre; MEK1^S217/221E^*mice have reduced levels of *Arc* mRNA (Xing et al., 2016). Hippocampal excitatory neurons in a developing forebrain-directed RASopathy mutant, *Emx1:Cre Ptpn11^D61Y^*, exhibit analogous reductions in depolarization-induced pERK1/2 and BDNF promoter activity (Altmüller et al., 2017). We propose the reduced levels of pERK1/2-ARC in cortical neuron soma and nuclei may be due to decreases in the development of cortical glutamatergic synapses or excitatory activity as a consequence of diminished long-range axon outgrowth.

In line with the reduced expression of *Arc*, our data show that *MEK1^S217/221E^* expression in *Nex:Cre* mice is sufficient to disrupt skilled motor learning. *Nex/Neurod6* predominantly drives Cre expression in immature cortical excitatory neurons, however, we cannot exclude that other *Nex/Neurod6* expressing neuronal populations could contribute to this behavioral phenotype (Goebbels et al., 2006). Nonetheless, restricting MEK1^S217/221E^ to layer V with *Rbp4:Cre* does not result in decreased skilled motor learning even though we observed a decrease in layer V neuron axon outgrowth. Motor learning is known to involve a complex, distributed set of neuronal populations that are not targeted in *Rbp4:Cre* mice. For example, the activity of layer II/III neurons in the motor cortex are recruited in a skilled reaching task and contain a sparse population of pyramidal neurons with some of the highest levels of pERK1/2 in the normal adult mouse cortex (Levy et al., 2020). We found a reduction in the number of layer II/III neurons with high levels of pERK1/2 in the motor cortex of *Nex:Cre; MEK1^S217/221E^* mice. Additional research is needed to explore the normal functions of these high pERK1/2-expressing layer II/III neurons in cortical processes and whether they are particularly vulnerable to RASopathic signaling.

## Acknowledgments

We wish to thank Noah Fry, Becca Reinking-Herd, and Sarah Bjorklund, for their technical assistance.

---

**Supplemental Figure 1.**
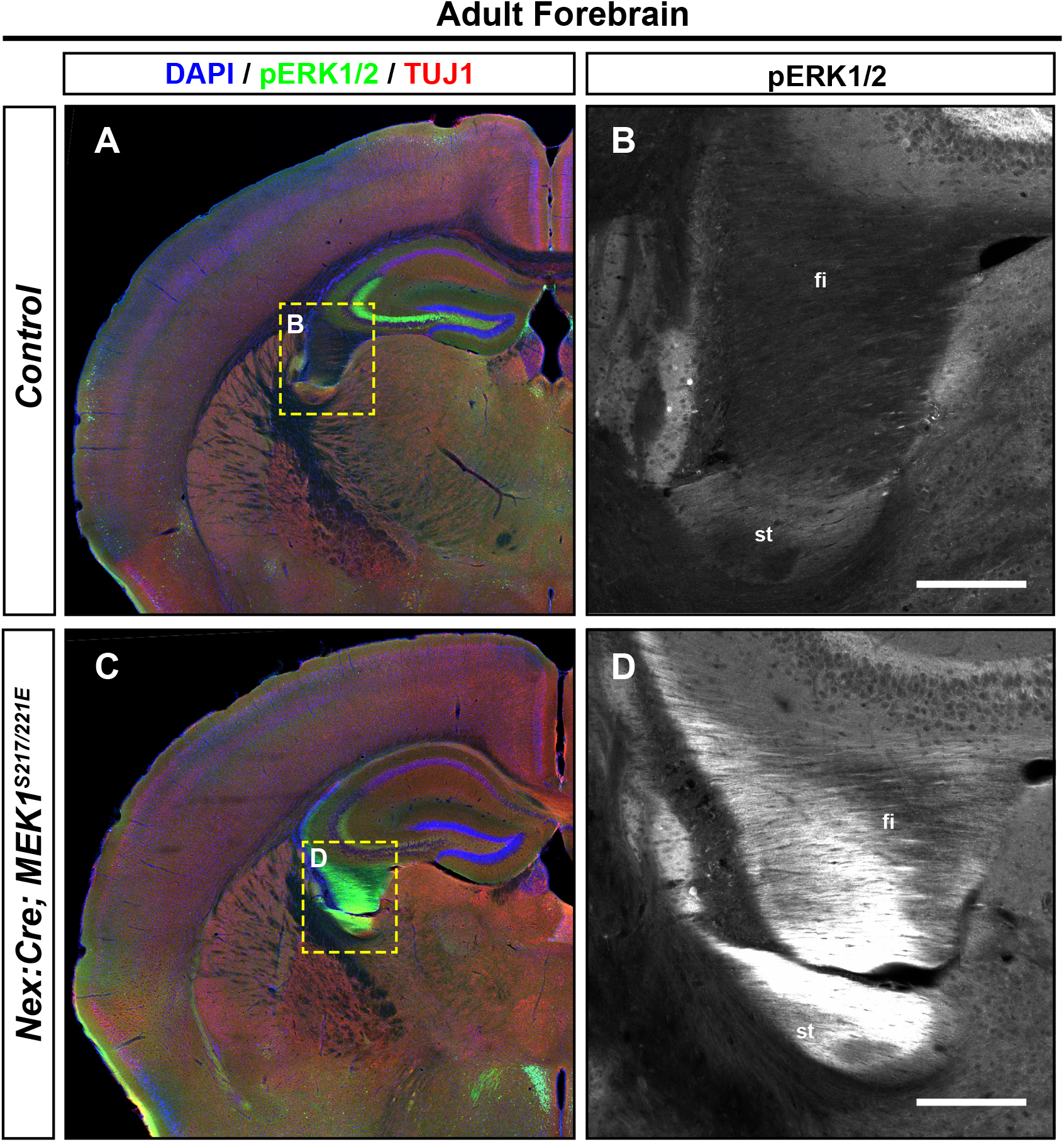
Increased levels of pERK1/2 in hippocampal axons of adult Nex:Cre; MEK1^S217/221E^ mice. **A-D.** Immunolabeling of forebrain sections at the level of the adult dorsal hippocampus showed a prominent increase of pERK1/2 within the mutant fimbria and stria terminalis (C-D) relative to controls (A-B), which are enriched in axons derived from excitatory pyramidal neurons in the hippocampal CA1-3 and the amygdala, respectively (n=4, fi = fimbria, st = stria terminalis). Scale bars = 200µm.

**Supplemental Figure 2.**
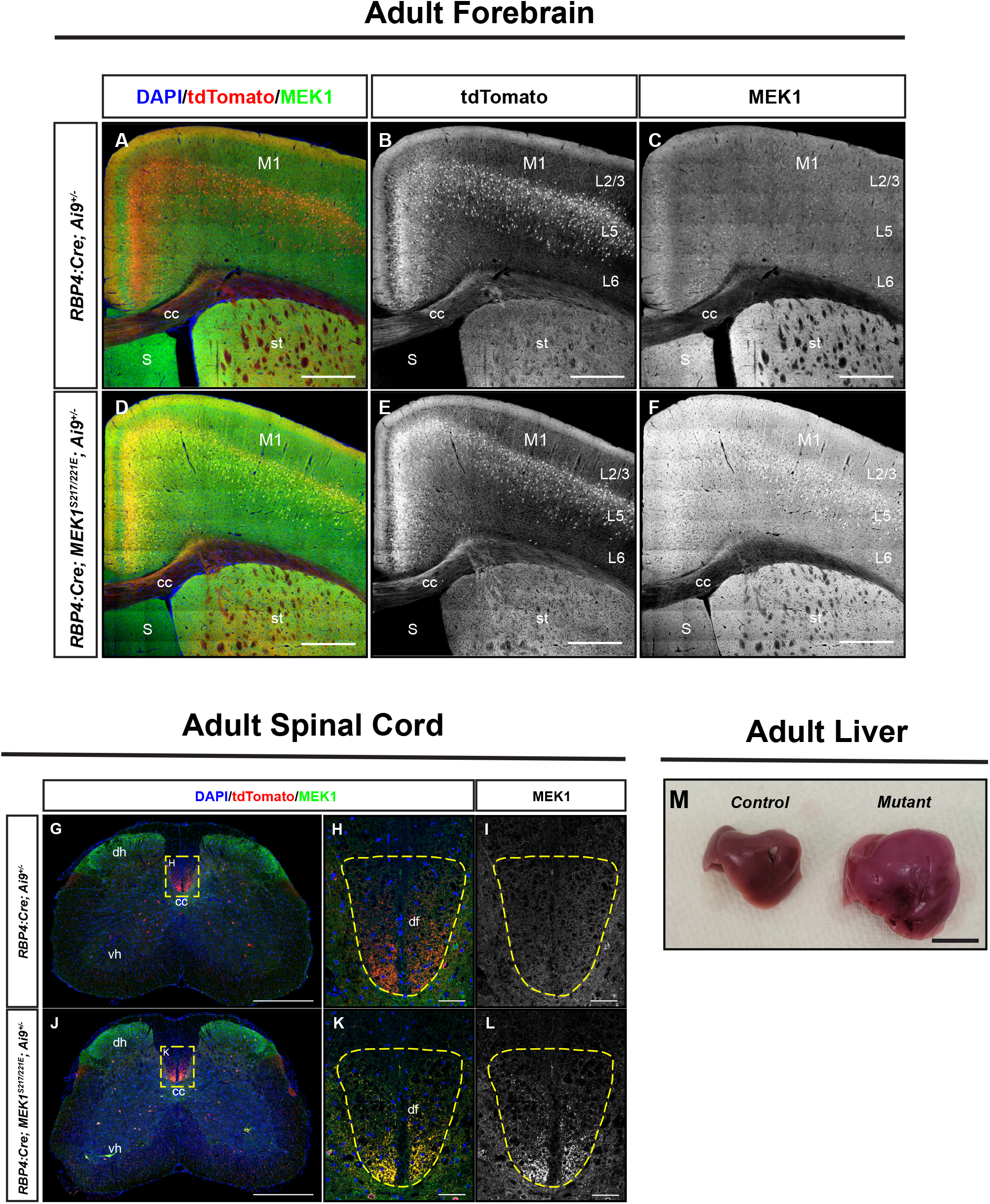
Conditional expression of MEK1^S217/221E^ in RBP4:Cre mice. **A-F**. Representative confocal images of *RBP4:Cre; Ai9^+/-^* cortices. Both *RBP4:Cre; Ai9^+/-^* mice control and *RBP4:Cre; MEK1^S217/221E^; Ai9^+/-^* mutants express RFP/tdTomato in cortical layer V (A-B, D-E). Immunolabeling of MEK1 showed a robust increase of expression restricted to cortical layer V in mutants (F) compared to controls (C). G-L. Representative images of cross-sectional lumbar segments showing that RFP is detected in descending projections in the dorsal funiculus of controls (G-H) and mutants (J-K). High resolution images reveal increased MEK1 expression in RFP-labeled axons of mutants (K-L) compared to controls (H-I). M. Gross dissection of the mutant liver was suggestive of hepatomegaly when compared to littermate controls (M). (M1 = primary motor cortex, cc = corpus collosum, st = striatum, S = septum, dh = dorsal horn, vh = ventral horn, cc = central canal, df = dorsal funiculus). Scale bars = Forebrain: A-F = 500 µm, Spinal Cord: G, K = 500µm, H-I, K-L = 50µm, Liver: M = 1cm.

**Supplemental Figure 3.**
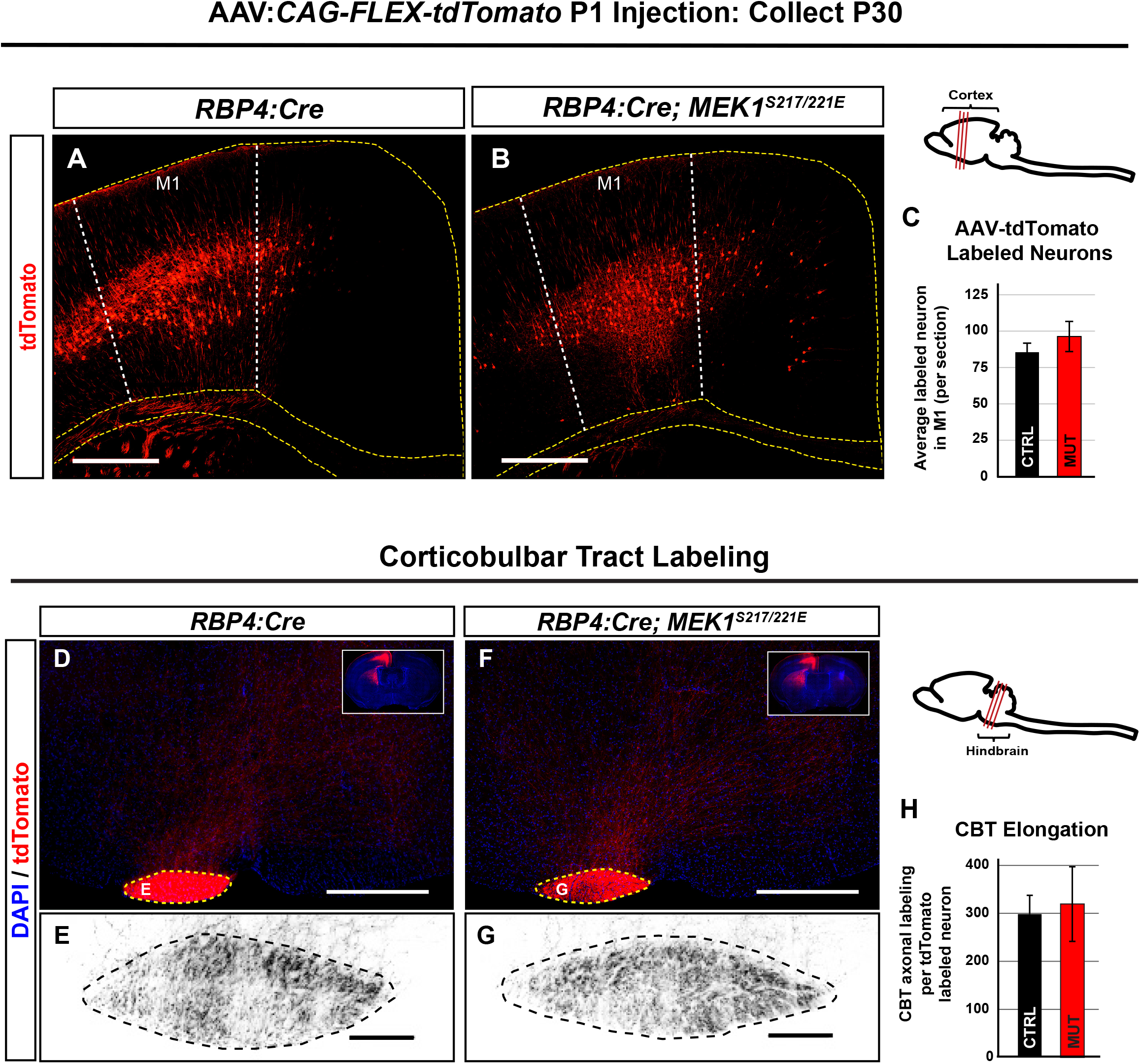
Corticobulbar tract axonal extension is not affected in *RBP4:Cre; MEK1^S217/2121E^* mice. **A-C**. Representative AAV injection sites in M1. AAV-tdTomato labeled cell counts in the motor cortex showed no significant differences between *RBP4:Cre;MEK1^S217/221E^* (B) and *RBP4:Cre* (A) mice as quantified in C (RFP^+^ cells per section, mean ± SEM, n=5, Student’s t-test p=0.4). **D-H**. Representative images of corticobulbar tract labeling. Quantification of tdTomato labeled axons in the corticobulbar tract revealed no significant differences between the *RBP4:Cre; MEK1^S217/221E^* (F-G) and *RBP4:Cre* (D-E) control mice (normalized to labeled neurons, mean ± SEM, n=4, Student’s t-test, p=0.818) (H). Scale bars: A,B,D,F = 500µm, E,G= 20µm.

**Supplemental Figure 4.**
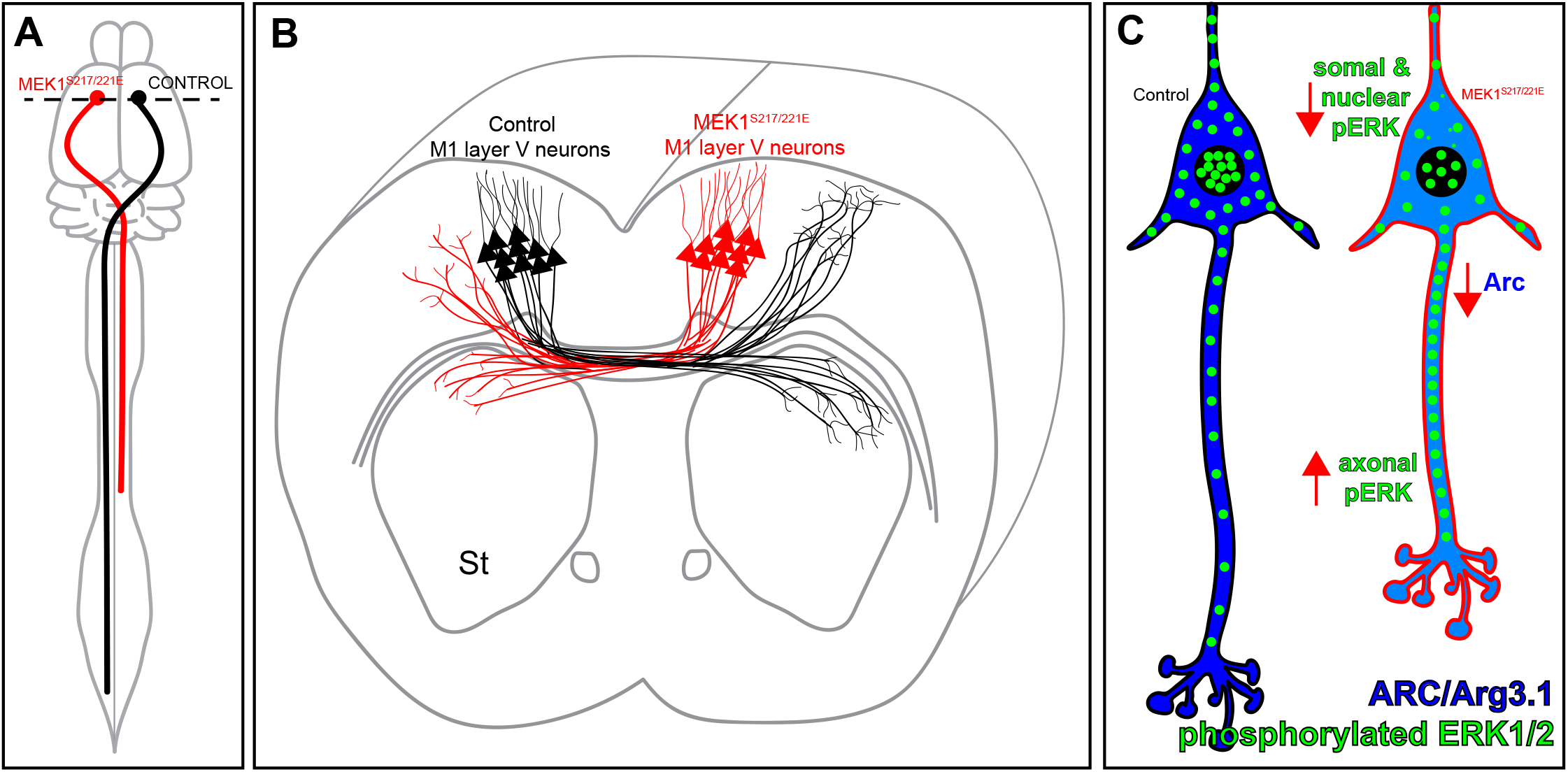
Model of aberrant circuit development in excitatory neuron-specific *MEK1^217/221E^* mice. **A-C**. *MEK1^S217/221E^* expressing layer V neurons (red) exhibit reductions in corticospinal axon elongation along the dorsal funiculus (A), reduced arborization in contralateral cortex and striatum (B), and persistent increases in axonal pERK1/2 (C - green dots) when compared to control neurons (black). These axonal deficits in *Nex:Cre; MEK1^S217/221E^* mutants coincide with decreased levels of pERK1/2 in the soma and nucleus and reduced expression of activity regulated cytoskeleton-associated protein, ARC (blue), which may underly select deficits in learning a skilled forelimb task (C).

